# Measurement of myofilament calcium in living cardiomyocytes using a targeted genetically encoded indicator

**DOI:** 10.1101/268003

**Authors:** Paul Robinson, Alexander J Sparrow, Connor N Broyles, Kolja Sievert, Yu-Fen Chang, Frances A Brook, Xiaoyu Zhang, Hugh Watkins, Yama A Abassi, Michael A Geeves, Charles Redwood, Matthew J Daniels

## Abstract

Visualising when and where calcium appears and disappears in cardiomyocytes is a major goal of cardiovascular research. Surprisingly we find that the chemical dyes widely used for this purpose disrupt cell contractility, due at least in part due to direct inhibition of the acto-myosin ATPase required to generate force. In order to improve calcium detection methods, we have developed a genetically encoded indicator that sits within the myofilament to directly visualise the changes occurring at the sarcomere. This tool improves on established chemical dyes and untargeted genetically encoded indicators for analysing small molecule modulators of myofilament-based calcium signalling. Importantly this is achieved without any measurable change in contractile function.

## Introduction

Calcium (Ca^++^) is the only physiologically occurring ion that activates contraction^1^. The process begins with voltage-dependent entry of extracellular Ca^++^ that is amplified by massive intracellular Ca^++^ release, principally from the sarcoplasmic reticulum (SR). The released Ca^++^ binds to Troponin C (TnC), promoting co-operative activation of the thin filament^2,3^ allowing the rigid actin scaffold to interact with the motor protein myosin enabling ATP dependent contraction^4,5^. Relaxation occurs as Ca^++^ is removed from the myofilament and repackaged into the SR, or pumped out of the cell, for subsequent cycles.

Protein architecture within the sarcomere is highly ordered and densely packed, producing a diffraction based striation pattern in unstained cells. Sarcomeric proteins are subject to point mutations and truncations in the common inherited heart diseases hypertrophic (HCM) and dilated (DCM) cardiomyopathy that change myofilament Ca^++^ sensitivity and/or contractility^6,7^. Modulators of contractility therefore represent potential therapies for these conditions, as well as the growing problem of heart failure^8^ in general. Importantly these should act without changes to total [Ca^++^] which is potentially pro-arrhythmic^9^. Therefore, to understand these diseases and identify such compounds, a need for myofilament [Ca^++^] and contractility assessment exists where ideally; measurement of one parameter does not alter the other.

Chemical dyes based on the non-fluorescent Ca^++^ binder BAPTA such as fura2^10^ or its colour variants are widely used to measure intracellular [Ca^++^] in cardiomyocytes. These are typically introduced into cells as acetoxymethyl (AM) esters, which increase membrane permeability and, after hydrolysis of the ester, lead to accumulation and concentration of the dye in the cardiomyocyte approximately 1000x the amount present in the medium. Adverse effects of these indicators have previously attributed to direct Ca^++^ buffering and phototoxicity. However, recent work has highlighted direct inhibition of the Na/K ATPase in many cell types by the BAPTA family of Ca^++^ indicators^11^ which limits cell survival and may reduce ATP dependent force production. Indeed fura2-based impairment of contractility has previously been observed in isolated rat cardiac trabeculae^12^.

Genetically encoded calcium indicators (GECIs) offer an alternative strategy for [Ca^++^] assessment^13^, with the possibility of subcellular targeting and lower (~μM) expression levels. However, their uptake in cardiovascular research has been slow^13^ compared to neurobiology^14,15^ as primary cardiomyocytes are difficult to transfect, and tend not to survive long enough in culture for gene expression and chromophore maturation. One highly desirable location for a Ca^++^ indicator is in the sarcomere, where calcium regulates contractility. Myofilament specific probes however remain elusive, in part due to the concern that the bulky 30×40×70Å adducts of the smallest GECIs^16^ may be poorly tolerated by the paracrystaline sarcomeric environment^7,17^ leading to perturbed function of the indicator, the fusion partner, or the cell.

Here we document the adverse effects on contractility of a number of chemical dyes widely used in cardiovascular research and go on to describe production and validation of a thin filament restricted GECI, RGECO-TnT. This novel Ca^++^ sensor, not only avoids any disruption of contractility, but also provides additional mechanistic information about Ca^++^ handling in the myofilament compared to an unrestricted RGECO when drugs that target the myofilament are used.

## Results

In a photometric system designed to minimise dye use and light exposure, we found the use of fura2 to measure [Ca^++^] in isolated ventricular adult guinea pig cardiomyocytes (vGPCMs) caused progressive inhibition of contractility across the dose range used in imaging studies (Figure 1a, b, **Supplementary Figure 1**). Impairment of fractional shortening may be caused by direct Ca^++^ chelation (fura2 *K*_d_ =145nM); as although the external concentration of fura2-AM ester is 1 μM, after ester hydrolysis fura2 becomes polar and cannot diffuse out of the cell leading to accumulation. Concentrations ranging between 0.1 and 40mM have been reported within cells^18,19^. Surprisingly, we also find inhibitory effects on whole-cell contractility with a chemical indicator of voltage, fluovolt^20^, and the sodium indicator SBFI^21^ (Figure 1c,d,e,f). These dyes do not bind Ca^++^ but do require weak surfactant application with or without acetoxymethyl esterification, and thus these data suggest there may also be Ca^++^ independent inhibitory effects of chemical dyes.

**Figure 1.**
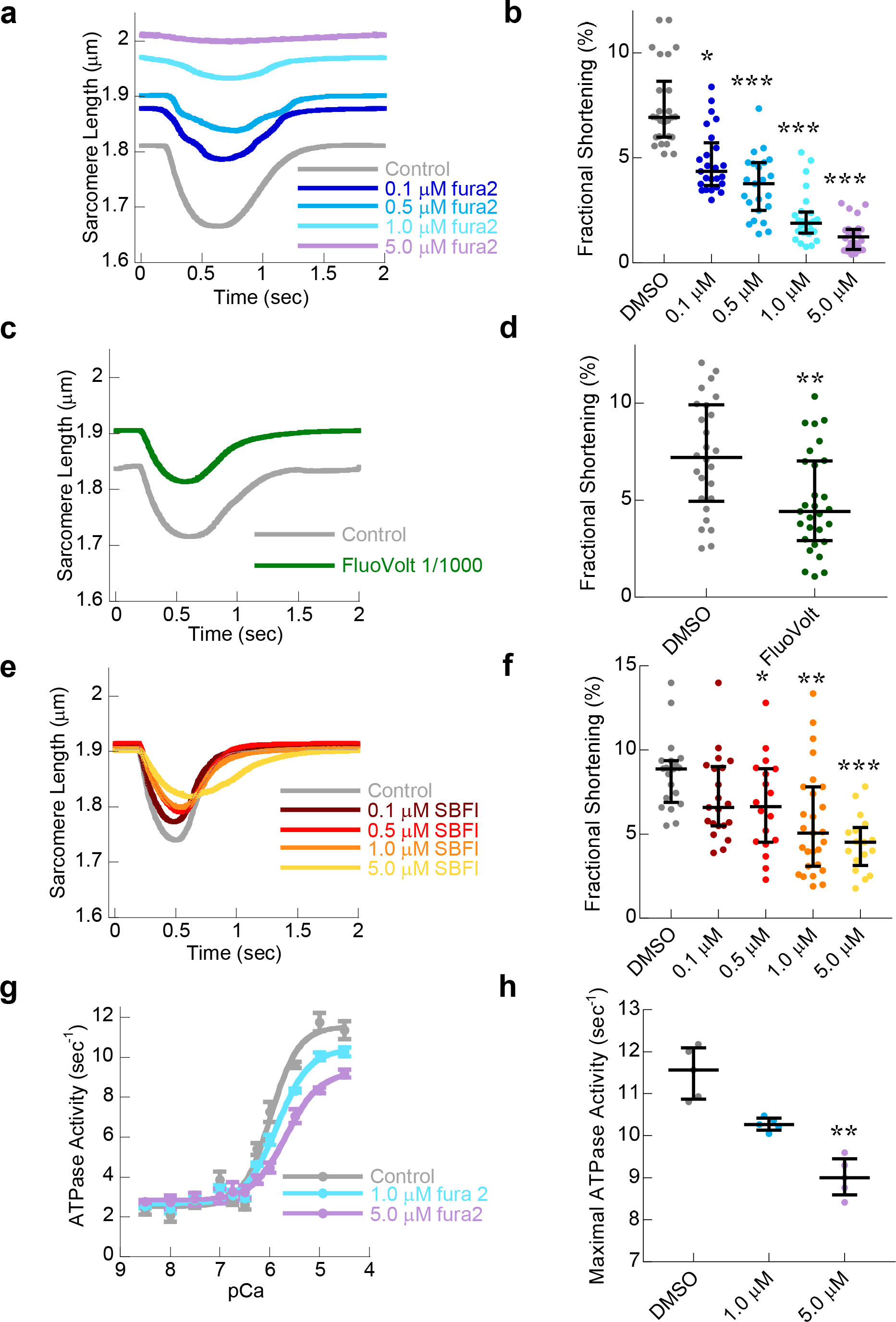
Chemical dyes to measure intracellular ionic flux is deleterious to cardiac contractility. Unloaded sarcomere shortening of electrically paced (0.5Hz) isolated adult cardiomyocytes was used to test cardiac contractile function in **(a,c,e)**. Control (grey) was compared to cells loaded with 0.1μM (royal blue), 0.5μM (blue), 1.0μM (teal) or 5.0μM (mauve) fura2-AM ester **(a)**. or with 1/1000 dilution of Fluovolt-AM ester (green) loaded according to the manufacturers instructions **(c)** or with 0.1μM (brown), 0.5μM (red), 1.0μM (orange) or 5.0μM (yellow) SBFI-AM ester **(e)**. Significant reduction in fractional shortening was observed in each case **(b)**. Working concentrations of Fluovolt-AM ester reduced contractile magnitude and caused basal relaxation of cardiomyocytes **(d)**. Exposure to increasing concentrations of the sodium dye SBFI-AM ester also reduced contractile magnitude however did not affect basal relaxation of cardiomyocytes **(f)**. Myofilament function was assessed using *in vitro* actin activated acto-myosin S1 ATPase assays. Myofilaments containing wild type control Tn complexes in DMSO (grey) were compared to those in the presence of 1μM (blue) or 5μM (mauve) fura2-AM ester. Note the shift of the trace down, indicative of myosin ATPase inhibition, as well as right **(g)**. n=5 error bars are ±SEM. The presence of fura2-AM ester reduced the maximal ATPase activity even with concentrations a fraction of the levels typically found in cells **(h)**. For all dot plots **(b,d,f,h)** lines give the median average and error bars are ± interquartile range * = p<0.05, ** = p<0.01 and *** = p<0.001 using one way ANOVA)

In light of recent data showing direct inhibition of the Na/K ATPase in many cell types by the BAPTA family of Ca^++^ indicators^11^ we investigated the Ca^++^ sensitivity of the actomyosin ATPase in vitro in the presence of fura2 to determine if Ca^++^ sequestration or ATPase inhibition was the cause of contractile impairment. In addition to reduced Ca^++^ sensitivity, the maximum rate of ATP hydrolysis is also significantly depressed suggesting direct inhibition of the actomyosin ATPase itself (Figure 1g,h) raising the possibility that other ATPases aside from Na/K ATPase may also be vulnerable to chemical dyes. Therefore, mechanisms beyond simple Ca^++^ buffering appear responsible for contractile impairment in dyes that do and do not have Ca^++^ affinity. To test the additional role of phototoxicity we made use of the adherent properties of iPS-derived cardiomyocytes (iPS-CM), which also show a dose response inhibition of the number of cells able to visibly contract in the presence of fura2 (**Supplementary Figure 2**) limiting the utility of motion tracking contractility assessments^22^ with simultaneous fura2 Ca^++^ imaging. Observer-independent impedance assessments of iPS-CM contractility can be undertaken without extrinsic illumination^23^. We find that contractile suppression is detected in the presence of green-shifted calcium indicators on monolayer iPS-CMs from a different vendor (**Supplementary Figure 3**); suggesting that neither UV light excitation phototoxicity occurring during fura2 imaging, nor cell singularisation, or indeed particular dyes or cell suppliers are required for this phenomenon. This suggests instead, that it may be due to direct biochemical effects of this group of compounds.

Protein based calcium indicators, the GECI’s, have been iteratively developed alongside improvements in fluorescent proteins over the past two decades^24^. Currently in cardiovascular research we lack probes that function where calcium is stored (the SR) and its major functional target (the myofilament). In order to develop a thin filament-restricted GECI we screened fusion proteins of green fluorescent protein (GFP) with various troponin subunits to identify sites that preserve actomyosin ATPase regulation. In contrast to the other fusion proteins tested, we found that GFP conjugated to the N-terminus of Troponin-T does not disrupt Ca^++^ regulation of the actomyosin ATPase (**Supplementary Figure 4, Supplementary Table 1**). Following this initial screen, the red calcium indicator RGECO^25^, previously applied to vGPCMs and iPS-CMs^26^, was conjugated to the N-terminus of Troponin-T (Figure 2a) and recombinant RGECO-TnT tested to show that it did not disrupt *in vitro* thin filament reconstitution (**Supplementary Figure 5)** or Ca^++^ sensitivity of the actomyosin ATPase (Figure 2b). In vitro characterisation of the calcium indicator properties of RGECO-TnT showed a *K*_d_ of 764.5±17.6 nM (RGECO was 860.0±10.0 nM) under physiological pH^27^, temperature and magnesium concentration^28^ (Figure 2c, **Supplementary Figure 6**), with preservation of dynamic range and excitation/emission spectra (Figure 2d and e, **Supplementary Table 2**). At physiological temperature, Ca^++^ on (at 10 μM free Ca^++^ *k*_obs_ 16.7 μM^−1^s^−1^) and off (*k*_off_ =16.9 s^−1^) rates for RGECO-TnT were similar to RGECO (**Supplementary Figure 7**), and approximately twice those observed at 25ºC.

**Figure 2.**
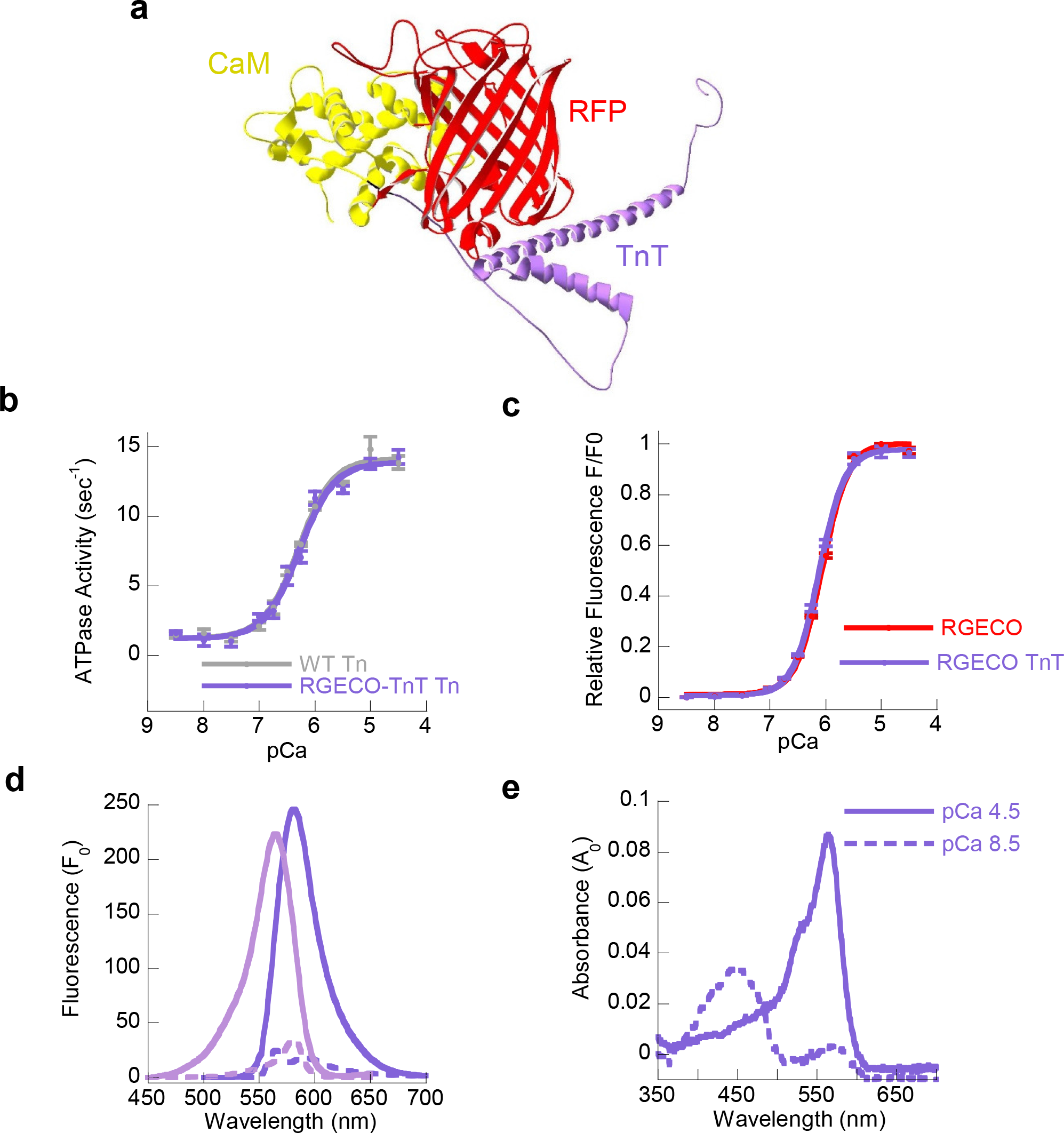
Conjugation of RGECO to the human cardiac TnT does not affect the function of the myofilament, or the indicator. Raptor X predicted structures from full length sequence **(Supplementary Figure 13)** submitted, returns all amino acid sequence of RGECO conjugated to amino acids 1-10 and 187-288 of human cardiac troponin T including the Gly-Ser linker. This indicates the C terminal T2 head thought to be essential for Tn reconstitution was freely available for interaction with the remaining Tn subunits **(a).** Myofilament function was assessed using *in vitro* actin activated acto-myosin S1 ATPase assays in **(b)**. The Ca^++^ sensitivity of control (unconjugated) troponin complexes (grey lines) were not significantly different to thin filaments reconstituted with RGECO-TnT (purple lines) in paired experiments. n=5 error bars are ±SEM. The steady state Ca^++^ binding affinity (K_d_) of purified recombinant RGECO (red lines) was compared to RGECO-TnT (purple lines) was assessed by analysis of the fluorescence/pCa relationship **(c)**. There is no significant difference in the fluorescence/pCa relationship between the two GECIs (n=4), error bars are ± SEM. Steady state fluorescence excitation, emission (peak excitation (Ex) = 564nm) / peak emission (Em) = 581nm) and absorbance spectra were obtained at pCa8.5 (dotted lines) and 4.5 (solid lines) for purified recombinant RGECO-TnT **(d and e)**. All experiments were performed at 37°C, 1.3mM MgCl_2_ pH7.3. Resultant molar extinction co-efficient, quantum yield, brightness and dynamic range measurements are given in **Supplementary Table 2**.

When RGECO-TnT is adenovirally expressed in vGPCMs (Figure 3a) or iPS-CMs (Figure 3b), in contrast to an unrestricted RGECO^26^, sarcomeric localisation without M line or Z-disc accumulation is observed. RGECO-TnT accumulates at levels similar to endogenous TnT (Figure 3c). With subcellular fractionation, we find that RGECO TnT shows exclusive accumulation at the sarcomere and is undetectable in the cytoplasm of vGPCMs (**Supplementary Figure 8**). Unlike fura2, which relaxes the resting sarcomere length and reduces sarcomeric shortening, the contractile parameters of both vGPCMs (Figure 3d & e) and iPS-CMs (**Supplementary Figure 2**) appear unaffected by the presence of either RGECO or RGECO-TnT. Ca^++^ transients are readily detected in electrically paced vGPCMs (Figure 3f) and iPS-CMs (Figure 3g), with reverse rate dependence. Dynamic range compression is apparent at higher pacing frequencies in vGPCM (**Supplementary Figure 9**) and iPS-CM (**Supplementary Figure 10**) as expected^26,29^. Spontaneous Ca^++^ transients are also observed in iPS-CM (**Supplementary movie 1**). GECIs can be employed in combination with other indicators^13,25^ or optical control tools^16,26^; indeed RGECO-TnT can be used simultaneously with the cytoplasmic green Ca^++^ indicator, GGECO^25^ in iPS-CMs (**Supplementary Figure 11**).

**Figure 3.**
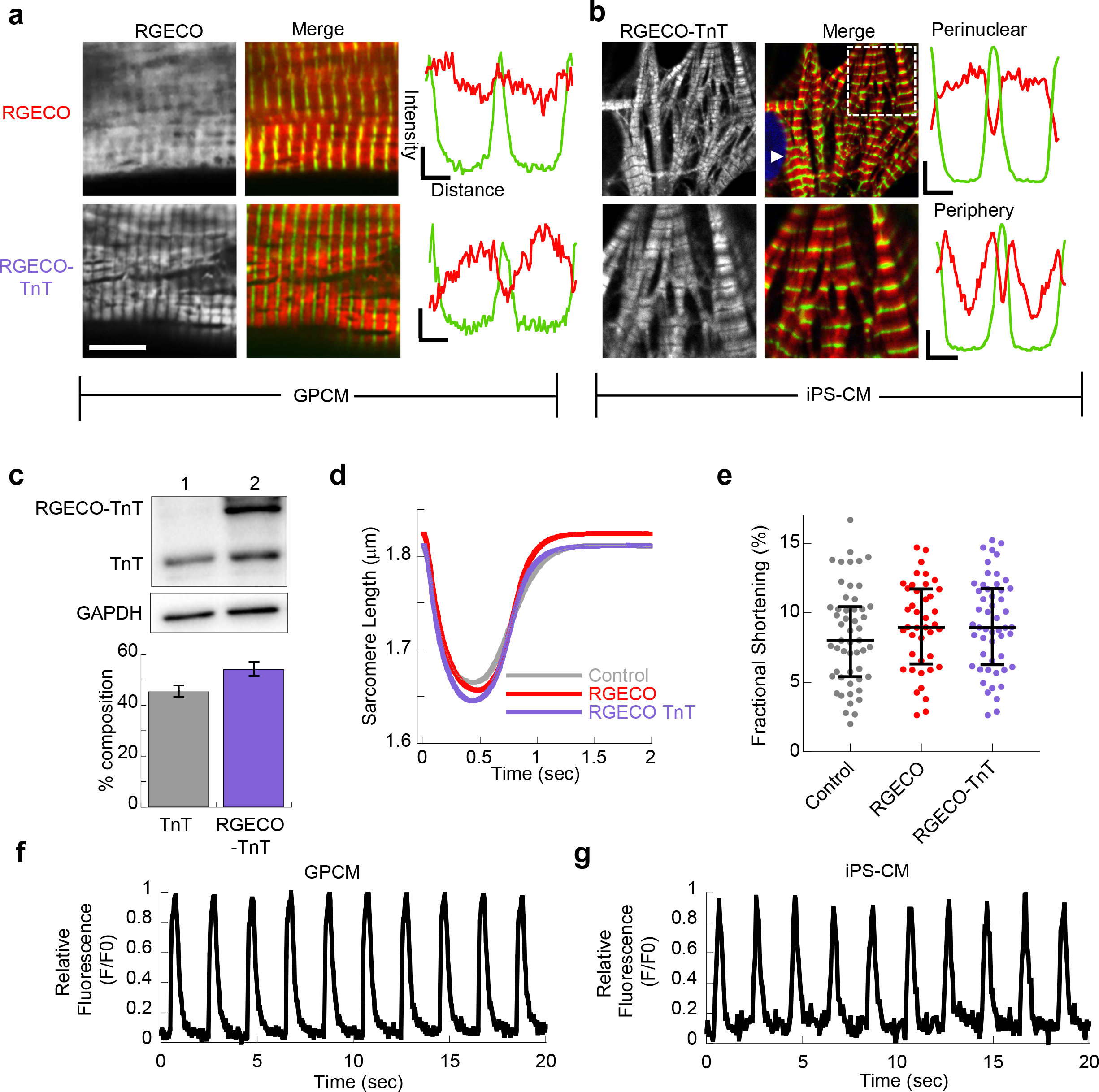
The red genetically encoded Ca^++^ sensor RGECO can be delivered to the myofilament by Troponin-T fusion revealing a dynamic signal from the sarcomere of adult ventricular and iPS derived cardiomyocytes without affecting contractility. In contrast to the diffuse staining of RGECO, adenovirally expressed RGECO-TnT localises to the I band in confocal microscopy of fixed adult ventricular **(a)** and human iPS derived cardiomyocytes **(b)**; z disks are revealed by α-actinin staining which is green in the merge image. Scale bar = 10μm. Dashed box shows the region expanded in the zoom panel. Intensity profile plots spanning two sarcomeres (three z-discs, ~3.6µm) are shown adjacent to the merge image, α-actinin (green line) labels the z-disc, the Ca^++^ indicator (red) is excluded from the Z-disc by TnT fusion. iPS-CMs appear to have different sarcomere morphologies adjacent to the nucleus (white arrow head), or the periphery (zoom panel). Western blot analysis of vGPCMs expressing RGECO-TnT in **(c)**, indicates that 54.3±5.5% (n=5) of endogenous TnT is replaced. Unloaded sarcomere shortening during electrical paced (0.5Hz) of isolated adult cardiomyocytes was used to test cardiac contractile function in vGPCMs by sarcomeric length measurement during contraction **(d)** or by fractional shortening assessment **(e)**. RGECO (Red) and RGECO-TnT (Purple) infected cells were not significantly different to control (Grey) **(e)**, error bars are ± interquartile range. The trace recordings of change in red fluorescence Ca^++^signal acquired from RGECO-TnT expressing single vGPCMs **(f)** and iPS-CMs **(g)** imaged at video rate during 0.5Hz electrical pacing.

There is growing interest in the therapeutic use of small molecule modulators of myofilament contractility^7,9,30,31^. Importantly these should not affect Ca^++^ handling, which may be pro-arrhythmic. To contrast the sensitivities and mechanistic insights of RGECO-TnT, RGECO and fura2, we investigated the effects of three drugs on Ca^++^ handling and contractility. The myosin ATPase inhibitor MYK-461^30^, levosimendan^31^ (a myofilament activator that stabilises the Ca^++^ bound form of TnC) and flecainide^26^, which does not target the myofilament but has complex effects on sodium (SCN5a)^32^, and potassium (hERG)^33^ channels, and the ryanodine receptor (RyR2)^34,35^ (Figure 4a, **Supplementary Tables 3a, 4a, 5a**). We find that the readouts of these probes are numerically and statistically different both in terms of their relative amplitudes and their timings. For example, MYK-461 reduces the relative fluorescence amplitude of RGECO (0.852±0.035, p=0.024) and more significantly using RGECO-TnT (0.802±0.037 p=<0.0004) whereas no change is seen with fura2 (0.913±0.056 p=0.124). RGECO-TnT also detects alterations to the T_50_ of Ca^++^ binding for all drugs tested (MYK-461 = −0.031±0.002 sec p=<0.0001, Levisomedan = −0.016±0.002 p=0.0001, Flecanide = −0.008±0.002 p=0.047) whereas both RGECO and fura2 show no significant differences of this parameter. Importantly the GECI probes are free from the inhibition of the actomyosin ATPase shown by fura2, and allows contractile responses equivalent to unlabelled cells (**Supplementary Figure 12, Supplementary Tables 3b, 4b, 5b)**, although these proteins will inevitably buffer intracellular Ca^++^. This remains true even when the indicator is concentrated in the myofilament itself where Ca^++^ regulates contractility.

**Figure 4.**
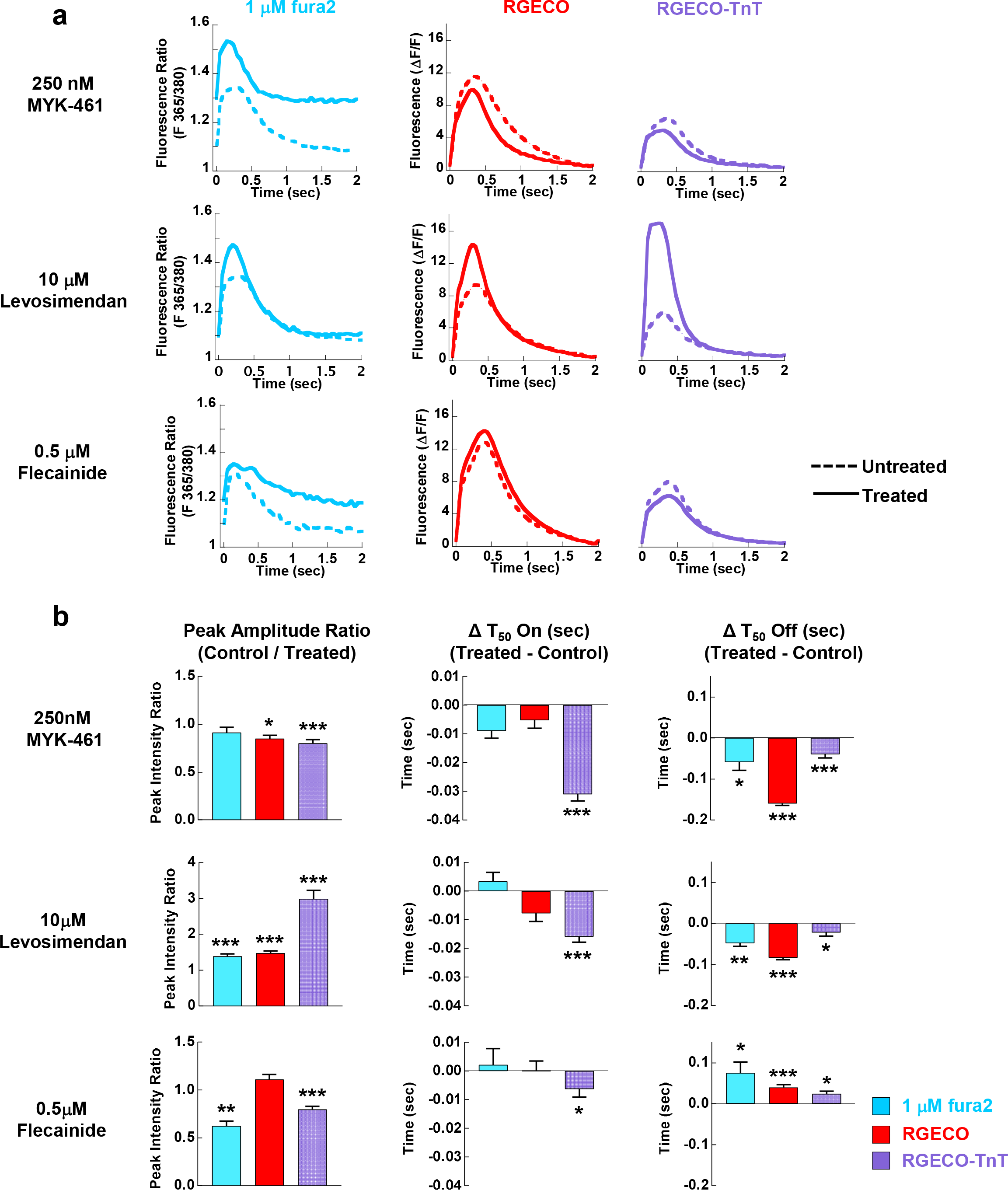
Small molecule effects on paired Ca^++^ transient measurements in vGPCM’s are different when fura2 is used compared to a genetically encoded probe RGECO or RGECO-TnT. The effects of MYK-461 (a myosin-ATPase inhibitor that reduces myofilament Ca^++^ sensitivity), Levosimendan (a myofilament Ca^++^ sensitizer) and Flecainide (a sodium channel inhibitor with off target effects on the potassium channel hERG, and the Ryanodine receptor RyR2) on the Ca^++^ transient and contractility were explored in electrically paced (0.5Hz) adult cardiomyocytes. The Ca^++^ dependent dynamic signals of 1μM fura2 (F365/380), RGECO and RGECO-TnT are shown in **(a)**. Each comparison is made in paired experiments between drug treated (solid lines) and DMSO control treated (dashed lines) for fura2 loaded (blue), RGECO infected (red) and RGECO-TnT infected (purple) cells. Δ values for all extracted parameters (including SEM and p values) for MYK-461, Levosimendan and Flecainide are plotted in **(b)**. Error bars are SEM, * = p<0.05, ** = p<0.01 and *** = p<0.001 using students t-test comparing untreated to treated cells. Absolute numbers for extracted parameters for MYK-461, Levosimendan and Flecainide treatment are tabulated in **Supplementary Tables 3, 4 and 5** respectively.

## Discussion

Chemical dyes are widely used to highlight the activation and inactivation of excitable cells on millisecond timescales. There are more than 25,600 citations for the use of BAPTA derivative Ca^++^ indicators to date^11^ but although problems like contractile inhibition^12^ and limited viability have hindered the field these effects were attributed to inevitable factors like calcium buffering and phototoxicity. Here we show that these mechanisms are less plausible than direct inhibition of the myosin ATPase itself, and in combination with a recent report documenting inhibition of the Na/K ATPase^11^ this raises the possibility that these compounds may have a general action as ATPase inhibitors. By contrast, in spite of their mechanism of action as Ca^++^ chelators GECI’s do not have biochemically detectable effects on the actomyosin ATPase *in vitro*, or on contractility at the whole cell level even when positioned inside the myofilament. This is in agreement with a recent report of a cAMP indicator fused to the C-terminus of troponin I (TnI) that gave correct myofilament localisation in cells and did not affect contractility^36^. We also tested the in vitro myofilament ATPase function of TnI as a smaller GFP fusion and found that typical Ca^++^ dependence was lost so this tool was not developed further. The discrepancy between these observations may be explained by relative expression levels of the cAMP reporter in adult rat cardiomyocytes, which we assume (as this was not detailed in^36^) will be lower than the 100 % used in our in vitro assay. Alternatively, the larger FRET reporter may exert less steric hindrance on troponin complex formation and native ATPase activity than the GFP / GECIs used in this study.

Myofilament targeting appears to enhance reporter sensitivity to the underlying biology. For example, RGECO-TnT detects larger peak amplitude reductions or increases compared to the other indicators strategies when small molecules that act via the myofilament (MYK-461 and levosimendan) are used. This may result from subcellular proximity of the sensor to site of drug action. The effect of MYK-461 on Ca^++^ handling was unexpected. Previous reports using fura2 measurements in isolated rat cardiomyocytes showed no effect on Ca^++^ transients with similar drug concentrations^30^. Although one explanation for this is the choice of indicator, other differences like duration of fura2 exposure, or different electrophysiology and Ca^++^ handling of rat cardiomyocytes compared to guinea pig cannot be excluded. When intracellular Ca^++^ was modulated with flecainide, the unrestricted GECI sees no change in signal amplitude, in contrast the myofilament restricted GECI sees it fall. Together these data suggest that RGECO-TnT improves the sensitivity of detection of Ca^++^ handling at the level of the myofilament when compared to fura2 or an unrestricted RGECO. These treatments also illustrate how the RGECO-TnT adds mechanistic information when assessing intracellular Ca^++^ dynamics. Whilst all three sensors detect changes to T_50_ off times, neither fura2 nor unrestricted RGECO reported changes to T_50_ on rates. However, the myofilament restricted sensor appears able to identify these subtle changes.

Both GECIs offer significant advances during combined assessments of contractility. While the presence of GECIs had no significant effect on sarcomere shortening, fura2 loaded cells relax and contract less and demonstrate composite effects due to the action of both fura2 and drug being tested. For example, in indicator-free control vGPCMs, 250nM MYK-461 increases resting sarcomere length by ~0.1μm. Either GECI strategy identifies this, but paradoxically the resting length shortens in cells also loaded with fura2. Similarly, although levosimendan treatment increases the resting diastolic sarcomere length in control cells, in the presence of fura2 it apparently decreases. These observations reveal the vulnerability for synergistic, or opposing, actions of the chemical dye on contractility to accentuate, or ameliorate, phenotypes induced by drugs in single cells. This may explain some of the difficulty generating these reagents, or their successful translation to the clinical environment.

In summary, we have identified a site within the contractile machinery that tolerates delivery of genetically encoded reporters into the heart of the myofilament without compromising the function of the cell, the indicator, or the fusion partner. This allows detection of the [Ca^++^] in the vicinity of the myofilament and permits simultaneous contractility measurements unaffected by the presence of the sensor, thus representing an improvement on the chemical dyes widely used in this field. This approach should also allow the targeting of other sensors to the myofilament to detect changes during development or disease.

## METHODS

### Adenoviral Virus design and production

Recombinant adenoviral constructs were created using the AdEasy XL recombination system (Agilent technologies) as previously described^37^. Briefly; RGECO and RGECO-TnT (**Supplementary Figure 13)** (sub-cloned using a BamH1 linker), were inserted into a shuttle vector, containing a CMV promoter. Viral DNA was produced via homologous recombination of the cloned shuttle vector, transfected and viral particles were scaled using the manufacturer’s standard protocol. Viral titre was estimated by infection of serial dilutions of purified virus in a plate containing ~2×10^6^ unmodified HEK293 cells over a 48 hour period. Cells were fixed using in 4% PFA and chemically skinned using 0.1% Triton X-100. Permeablized cells were incubated sequentially with a 1 in 200 dilution of Ds-Red primary antibody (Living Colors, Takara Bio, USA) and a 1 in 1000 dilution of Alexa 568 secondary antibody (Life Technologies, CA, USA). Subsequent cell counts of red fluorescent vs. total cell number gave a viable virus particle estimate of 5×10^10^/ml and 1.49×10^9^/ml for RGECO and RGECO-TnT respectively.

### Isolation of guinea pig left ventricular cardiomyocytes

This investigation was approved by the Animal Welfare and Ethical Review Board at the University of Oxford and conforms to the UK Animals (Scientific Procedures) Act, 1986, incorporating Directive 2010/63/EU of the European Parliament. A 400g male guinea pig (Envigo, Bicester, UK) was dispatched by cervical dislocation. Cardiomyocytes were dissociated using Langendorff perfusion of, 0.8mg/ml collagenase type II (Worthington Biochemical Corporation, Lakewood, NJ, USA) in isolation solution containing 130mM NaCl, 23mM 4-(2-hydroxyethyl)-1-piperazineethanesulphonic acid (HEPES), 21mM glucose, 20mM taurine, 5mM creatine, 5mM MgCl_2_, 5mM Na pyruvate, 4.5mM KCl, 1mM NaH_2_PO_3_, pH7.3 with NaOH). Cells were resuspended and plated at ~1×10^5^ per ml in ACCITT3 cardiomyocyte culture medium as first described by Ellingsen et al^38^ containing 500 ml M199 media 100 units of penicillin, 0.1 mg/ml streptomycin, 20mM taurine, 5mM creatine, 5mM Na pyruvate, 2mg/ml BSA, 2mM L-carnitine, 0.1pM insulin. Recombinant adenovirus was immediately added to 0.75ml of cell suspension to an estimated multiplicity of infection (MOI) of between 500 and 800, infected cells were placed at 37°C in a 5% CO_2_ atmosphere for 48 hours. This was sufficient to infect 100% of all viable rod shaped cardiomyocytes with either RGECO or RGECO-TnT.

### Stem cell derived cardiomyocytes

Human iPS derived cardiomyocytes and maintenance media were purchased from Axol Bioscience, (Cambridge, UK), and handled according to manufacturer’s recommendations at 37ºC, 5% CO2 in a humidified incubator. After 72 hours in culture they were singularised and re-plated onto Fibronectin (0.5%)/Gelatin (0.1%) coated 35mm glass bottom dish (No. 0) MatTek plates (Ashland, MA, US) at 1:6.

### Protein purification

N and C terminal eGFP conjugates of human recombinant TnT and TnC were produced by molecular cloning in PMW172 vector using flanking 3’ NdeI and 5’ EcoRI restriction sites and an internal HindIII restriction site used as a linker in each case (NEB, New England). RGECO and RGECO-TnT were also sub-cloned in to PMW172 using the same strategy. Proteins were expressed in BL21-DE3-pLysS *E.coli* induced using 0.4mM Isopropyl β-D-1-thiogalactopyranoside for 4 hours. Bacterial pellets were recovered by centrifugation (10,000 x*g* for 10 mins) and lysed in a buffer containing 25mM Tris HCl, pH 7.5, 20% sucrose, 1mM EDTA, 200mM NaCl, 5M urea, 0.1% Triton X-100. All proteins were purified using an AKTA-UPC900 FPLC using HiTrap FF chromatography columns (GE Healthcare, Amersham). GFP-TnT and TnT-GFP were purified using sequential cation (in buffer containing 6M urea, 1mM EDTA, 1mM 2-mercaptoethanol, 20mM MOPS, pH 6.0) followed by anion (in buffer containing 6M urea, 1mM EDTA, 1mM 2-mercaptoethanol 50mM Tris HCl pH8.0) exchange chromatography. GFP-TnC and TnC-GFP were purified by anion (pH8.5) exchange chromatography. TnI-GFP was purified by sequential cation and anion exchange chromatography. RGECO and RGECO-TnT were purified using sequential anion (pH8.0) exchange chromatography, Ammonium Sulphate fractionation (to 50% for RGECO and 35% for RGECO-TnT) and finally Hydrophobic Interaction Chromatography (HIC) (in buffer containing 30% ammonium sulphate, 200mM NaCl_2_ 1mM ditiothretol and 30mM 4-(2-hydroxyethyl)-1-piperazineethanesulphonic acid (HEPES)). Ion exchange columns were eluted using a gradient of 0-2M NaCl, the HIC column was eluted with a gradient of 30–0% ammonium sulphate. All eluted protein fractions were assessed for purity using 12% SDS PAGE gels, stained with coomassie brilliant blue. Wild type human recombinant TnT, TnI, TnC and Ala-Ser-α-TM^39^ were purified as previously described^40^. Troponin complex containing either WT, GFP-TnT, GFP-TnT, GFP-TnC, TnC-GFP or RGECO-TnT were reconstituted by dialysis into buffer containing 10mM imidazole pH7.0, 1mM DTT, 0.01% azide, 0.1mM CaCl_2_, 6M urea and 1M KCl, firstly urea was reduced stepwise from to 2 then 0M then KCl was reduced stepwise from 1M to 800 – 600 – 400 - 200mM KCl in a series of 3 hour dialyses. Tn Complex was purified using size exclusion chromatography in 200mM KCl dialysis buffer (**Supplementary Figure 5a**), purity was analysed by SDS PAGE (**Supplementary Figure 5b**), and finally, purified troponin complexes are dialysed into buffer containing 5mM 1,4-Piperazinediethanesulphate pH7.0, 3,87mM MgCl_2_, 1mM DTT for ATPase assay experiments. Actin and myosin-S1 was extracted from rabbit skeletal muscle as previously described^41,42^.

### Actin Co-sedimentation assays

Reconstituted Tn complex proteins (0.5μM) were co-sedimented with human Ala-Ser-α-TM (0.5μM) and rabbit skeletal actin (3.5μM) at 384000x*g* for 15 mins by ultracentrifugation (Beckman Coulter Inc, High Wycombe). Total, supernatant and pellet fractions were visualized using coomassie blue stained 12% SDS PAGE gels in order to ensure complete reconstitution of thin filaments prior to all ATPase assay measurements. (**Supplementary Figure 5c**)

### *In vitro* acto-myosin activated myosin ATPase assays

ATPase assays were undertaken using our standard assay conditions^6,40^ using 3.5μM actin, 0.5μM myosin S1, 0.5μM Ala-Ser-α-TM and 0.5μM Tn complex in buffer containing 5mM 1,4-Piperazinediethanesulphate pH7.0, 3.87mM MgCl_2_, 1mM DTT. In order to ensure precise thin filament protein stoichiometry, each stock was centrifuged at 384000x*g* for 15 mins. Supernatants were discarded, actin pellets were recovered in buffer containing 5mM 1,4-Piperazinediethanesulphate pH7.0, 3.87mM MgCl_2_, 1mM DTT to an equal volume of the discarded supernatant. Reaction mixtures were aliquoted and set to a range of _free_[Ca^++^] between 3.16nM (pCa8.5) and 31.6μM (pCa4.5) using 1mM EGTA and the appropriate corresponding concentration of CaCl_2_, calculated using Maxchelatior software (http://maxchelator.stanford.edu/CaEGTA-TS.htm). Calcium-sensitivity data was fitted to the Hill equation using Kaleidagraph (Synergy Software, Inc (USA)).

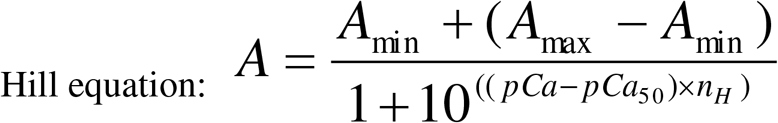

Where: A = ATPase rate; A_min_ = Minimum ATPase rate; A_max_ = Maximum ATPase rate; pCa = −log [Ca^++^]; pCa_50_ = - log [Ca^++^] required for half maximum ATPase activity; *n*_H_ = Hill coefficient.

### Calcium binding, *K*_d_ and kinetic calculations

To measure steady state Ca^++^ binding affinity (*K*_d_) for RGECO and RGECO-TnT, 3μM of protein was dialysed into buffer containing 130mM NaCl, 10mM 4-(2-hydroxyethyl)-1-piperazineethanesulphonic acid (HEPES), 1mM dithiothretol, pH7.3 and either 0, 1.3 or 3.9mM MgCl_2_. A range of free[Ca^++^] conditions were set between 3.16nM (pCa 8.5) and 31.6μM (pCa 4.5) using 1mM EGTA and the appropriate corresponding concentration of CaCl_2_. Steady state fluorescence readings were made in an Ultraclear bottom 96 well microplate using a FLUOstar Omega plate reader (BMG LABTEC, Ortenberg, Germany), using a 544nm excitation filter and 590/10nm emission filter at both 25 and 37°C. Resultant fluorescence emission intensities were plotted vs _free_[Ca^++^] and fitted to the Hill equation to calculate *K*_d_ values for each protein and condition.

To measure Ca^++^ displacement (*k*_off_), 125nM RGECO or RGECO-TnT plus 5µM CaCl_2_ was mixed with 5mM EGTA in a buffer containing 130mM NaCl, 10mM 4-(2-hydroxyethyl)-1-piperazineethanesulphonic acid (HEPES) 1.3mM MgCl_2_, 1mM dithiothretol, pH7.3 with NaOH. RGECO or RGECO-TnT was loaded into a Stopped-flow system (HiTech Scientific, Bradford-on-Avon, UK), concentrations after mixing 1:1 in the stopped-flow. Fluorescence was excited at 546nm (100W Xe/Hg lamp and monochromator) and emission monitored through a OG-590 glass filter. The resultant data was fitted to a single exponential decrease in fluorescence of 50-100%, depending upon temperature. To measure calcium binding (*k*_on_), 125nM RGECO or RGECO-TnT was measured at 10µM free calcium by mixing with a buffer containing 2.125mM Ca.EGTA and 0.2mM EGTA. The stopped-flow fluorescence filter set up was idetical to that of the displacement measurements. A large increase in fluorescence upon addtion of calcium was fitted to a single exponetial. The observed single exponential rate constant, *k*_obs_, was extracted for *k*_on_ and *k*_off_ for both sensors, at a range of temperatures between 25 and 37°C. To estimate the dissociation constant we using the equations *K*_d_ = *k*_obs_-off [Ca]/ *k*_obs_ on. Which were of comparable values to steady state *K*_d_ values obtained previously.

### Determination of Quantum Yield and Molar Extinction Co-efficient

Quantum yield and molar extinction co-efficient was determined for each Ca++ sensor using a method adapted from Zhao *et al*^25^. Briefly, standards for quantum yield determination were mCherry (for RGECO) and RGECO for RGECO-TnT. Briefly, the concentration of protein in buffer containing 10mM 4-(2-hydroxyethyl)-1-piperazineethanesulphonic acid (HEPES), 1.3mM MgCl_2_ and 1mM dithiothreitol was determined using BCA assay (Pierce) and set to either pCa 4.5 of 8.5, the concentration of each protein was reduced to an absorbance at the excitation wavelength between 0.2 and 0.6. A series of dilutions to absorbance’s of 0.01, 0.02, 0.03 and 0.04 were made for each protein solution and standard. The fluorescence emission spectra at Ex F_568_ of each dilution of each standard and protein solution were recorded using an RF-1501 spectrofluorimeter (Shimadzu, Kyoto, Japan). The total fluorescence intensities were then obtained by integration. Integrated fluorescence intensity vs. absorbance was plotted for each protein and each standard. Quantum yields were determined from the slopes (S) of each line using the equation: Φprotein = Φstandard × (Sprotein/Sstandard).

Extinction coefficients were determined by first measuring the absorption spectrum of both RGECO and RGECO-TnT at pCa8.5 and 4.5 in the buffer above using a plate reader (BMG LABTECH, Offenburg, Germany). Peak Absorbance wavelengths were determined to be A_465_ in low Ca^++^ and A_581_ in high Ca^++^. Absorbance measurements were automatically converted to 1 cm path length using BMG Omega software. Extinction coefficients of each protein were calculated by dividing the peak absorbance maximum by the previously determined concentration of protein.

### Subcellular fractionation and western blotting

Adult guinea pig left ventricular cardiomyocytes were fractionated to recover cytoplasmic and sarcomeric/cytoskeletal samples in a method adapted from Simon *et al^43^*. Briefly, 300,000 cardiomyocytes were pelleted by centrifugation, resuspended in buffer containing, 20mM Tris HCl pH7.4, 2mM EDTA, 0.5mM EGTA, 0.3mM sucrose. Cells were homogenised in a glass homogeniser for 1 minute. Cells were pelleted at 1000xg for 5mins, the supernatant / cytoplasmic fraction was retained for SDS PAGE analysis and western blotting. The cell pellet was washed 3 times in fractionation buffer + 1% Triton X-100 each time pelleting the sarcomeric fraction by centrifugation at 1000xg for 5mins. Finally, the sarcomeric fraction was resuspended in fractionation buffer (without Triton X-100) and prepared for SDS PAGE analysis and western blotting. Cytoplasmic and Sarcomeric fractions were western blotted using anti-cTnT (1 in 2000) (Sigma, Poole, UK) and anti-ERK (1 in 4000) (Cell Signaling Technologies, Massachusetts, USA) primary antibodies and rabbit HRP (1 in 8-16000) secondary antibodies using PVDF membranes and visualised and analysed using GelDoc Image Lab software (Bio-Rad Laboratories, California, USA).

### Measurement of sarcomere shortening and fura2 Ca^++^ transients

Sarcomere shortening and Ca^++^ transient measurements were performed using IonOptix µstep apparatus and the manufacturers’ standard operating instructions. Briefly: cultured cardiomyocytes were loaded with fura2 Ca^++^ indicator by incubation with either 0.1, 0.5, 1.0 or 5.0μM fura2-AM ester (Life Technologies) in the presence of 2M F127 pluronic in ‘perfusion buffer’ (150mM NaCl, 10mM 4-(2-hydroxyethyl)-1-piperazineethanesulphonic acid (HEPES), 7mM glucose, 1mM MgCl, 1mM KCl, 0.3mM NaH_2_PO_3_, pH 7.4 with NaOH) containing 250µM CaCl_2_ for 5mins, followed by a 10 minute wash in perfusion buffer containing 500µM CaCl_2_ to remove any excess label. For experiments using SBFI 0.1, 0.5, 1.0 or 5.0μM were loaded in the same manner. Experiments using FluoVolt (ThermoFisher) a 1/1000 dilution was used according the manufacturer’s instructions. The loaded cells were then allowed to settle to the bottom of a perfusion chamber with a 1.5 thickness cover slip base, which was mounted on an inverted fluorescence microscope. Cells were perfused with Perfusion buffer containing 1.8mM CaCl_2_, and electrically paced at 40 volts. Pacing frequency was set at 0.5Hz in order to accurately measure resting diastolic [Ca^++^]_i_. Sarcomere shortening was captured by Fourier transform of the cardiomyocyte striations under phase contrast microscopy using a switching rate of 100Hz. fura2 Ca^++^ transients were captured simultaneously, using the ratio of fura2 fluorescence emission at 365/380nm at a switching rate of 1000Hz. All contracting cardiomyocytes were measured for contractility and fura2 Ca^++^, any cells displaying asynchronous contractility, excessive blebbing/dysmorphology were ignored for acquisition, whilst any cell with contractile magnitudes or velocities exceeding 2 standard deviations from the mean upon analysis were also excluded as product of phenotypic heterogeneity of primary cells in culture losing function and morphology over time. To compare sarcomere shortening and fura2 Ca^++^ transients in the presence of contractility or Ca^++^ modifying small molecules MYK-461 (0.25μM), Levosimendan (10μM) and Flecainide (0.5μM), 200 μl of cultured (48 Hours) cardiomyocytes were incubated in either drug or DMSO vehicle for 5mins. The entire cell preparation was pipetted onto the microscope perfusion chamber buffer containing the either drug or vehicle was perfused at 0.75ml/min for the duration of the experiment. All contracting cells were measured (except those not meeting the exclusion criteria above), no preparation of cells was left for more than 15mins before being replaced for a fresh batch of pre-incubated cells. At least 3 cell preparations were analyzed for each DMSO/drug comparison. No differences or clustering was observed between each cell preparation assessed.

### RGECO imaging in fixed cells

Adult ventricular cardiomyocytes were exposed to recombinant adenovirus immediately after isolation in 0.75ml of cell suspension to an estimated multiplicity of infection (MOI) of between 500 and 800, infected cells were placed at 37°C in a 5% CO_2_ atmosphere for 48 hours. At 48 hours an aliquot was taken and paraformaldehyde to a final concentration of 4% was added, at 10mins cells were pelleted (600rpm, 2mins) in a benchtop centrifuge, washed with 1xPBS and resuspended in PBS. Cells were spun (Cytospin, Thermo Scientific) onto glass slides, and ringed with a PAP pen (Sigma). Human iPS-derived cardiomyocytes were plated as above and incubated to day 7 following thaw before washing with 1xPBS and fixation with 4% paraformaldehyde. For infection, virus was diluted 1:10 from the stock solution and used at 1μl/dish 24 hours after replating. Permeabilisation with 0.1% Triton X-100 in Tris Buffered Saline for 10mins at room temperature was followed by blocking (0.2% albumin in permeabilisation buffer) for 20mins. Primary antibodies (Mouse monoclonal anti-alpha actinin (Sigma, clone EA-53) rabbit polyclonal anti-DsRed (Clontech) were diluted 1:2000, and 1:200 in blocking buffer respectively. Three hours after primary incubation cells were washed in permeabilisation buffer, and counter stained with Alexa-488 anti-mouse, and Alexa-568 anti-rabbit fab fragment secondary antibodies (Invitrogen), nuclear counterstaining was with TO-PRO-3 (Invitrogen), for an hour before washing and mounting (Vectashield, Vector labs). Images were acquired on a Leica SP5 confocal microscope with a 63x oil immersion lens, and intensity plots prepared with the plot profile function in ImageJ.

### Live cell Imaging Microscope

An inverted IX81 frame (Olympus, Japan), with a custom 7 LED array (Cairn Research, Faversham, UK), automated stage (Prior Scientific, Cambridge, UK), lens turret, and filter wheel is housed in a custom heated, humidified chamber (Solent Scientific, Segensworth, UK) with image collection on two C-1900 EMCCD cameras (Hamamatsu, Japan) mounted on a DC2 beam splitter (Photometrics, UK). For isolated RGECO imaging, single camera mode image acquisition through an Olympus LUCPlanFLN 40x objective lens (NA 0.60), or an Olympus UPlanFLN 10x lens (NA 0.3), using CellR software (Olympus) was used with uninterrupted recording of the calcium transient via a 5msec 568nm LED illumination pulse and the RFP filter set (DS/FF01-560/25-25, T565lpxr dichroic mirror, and ET620/60 emission filter, all Semrock). Simultaneous calcium transients from GGECO and RGECO indicators was acquired through an Olympus PlanApo 60x oil objective lens (NA 1.42), through a quad-band filter set (band pass filter (DS/FF01-387/485/559/649-25), dichroic quad-edge beam splitter (DS/FF410/504/582/669-Di01-25×36), and quad-band emission filter (DS/FF01-440/521/607/700-25) all Semrock with fluorescent emissions recorded through a Dual-View system (DC2, Photometrics) with green (520/30nm) and red (630/50nm) channels to separate cameras controlled through CellR (Olympus, Japan). RGECO and RGECO-TnT adenovirally induced vGPCMs were treated with 250nM MYK-461, 10µM Levosimendan, or 0.5µM Flecainide for 20mins then video rate movies with 0.5Hz pacing were acquired for 28secs using the UPlanFLN 10x lens.

### Transient analysis software & statistics

Raw image data was extracted using CellR (Olympus), and analysed in Excel (Microsoft). From a single cell 10 intensity transients over time were extracted and averaged to give a single transient in a 2 second interval per cell; 3 transients were averaged for iPS-CMs. Time to 50% contraction, or time to 50% baseline (from peak) was determined from these traces.

Comparative peak intensity analysis was performed using paired control vs drug exposed datasets obtained from the same cell preparation. In brief the ΔF/F measurement described for intensiometric indicators^34^ was used.

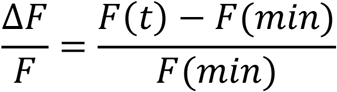

Where *F*(*t*) is the fluorescence intensity at time (t), and F(min) the minimum fluorescence intensity during the 2 second averaged dataset taken to be representative of a single cell.

Comparisons presented in Figure 2b were obtained by dividing or subtracting the relevant value of the paired control population from that of the drug exposed population based on the data in Supplementary tables 3, 4 & 5.

Where relative intensity data is presented, the following calculation was used

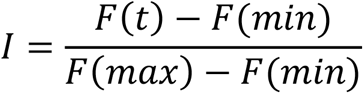

Where *I*, intensity, is a ratio of the fluorescence signal (*F(t)*) scaled according to the maximum (*F(max)*), and minimum (*F(min)*) response of the cell during the observation.

Unfiltered transient graphs were plotted using KaleidaGraph (Synergy Software), and dot plots with GraphPad Prism (GraphPad software). Groups were compared using unpaired Students t-test (MYK-461, Levosimendan, Flecainide) or one-way ANOVA with post-hoc Dunn’s test (pacing frequency comparisons). Simultaneous RGECO and GGECO calcium traces from a single cell, and average responses at different pacing frequencies were normalized for presentation.

### iPS-derived cardiomyocyte movement analysis

Cells plated and prepared as above were imaged in bright field mode at video rate over a 2 minute interval using the LUCPlanFLN 40x lens. Quantification of cell activity was scored for actively/non-actively contracting cells. An activity contracting cell was regarded as a cell presenting at least a single contraction within the 2 minute interval. The spontaneous beat frequency for these cells is 0.52Hz +/− 0.15Hz, with >90% cardiomyocyte purity based on TnI positivity^26^.

### Impedance assessment of contractility

iCell cardiomyocytes (CDI) were seeded at 20,000 cells/well into RTCA plates and loaded onto an xCELLigence RTCA Cardio system (ACEA bioscience) housed in a humidified incubator at 37°C with a 5% CO_2_ atmosphere, this was interrupted for feeding on alternate days. Contractility was monitored via impedance assessment at 12.9msec intervals. At day 14 post seeding, with robust signals apparent in all wells, cells were loaded with FLIPR 5 (Molecular Devices) around the recommended 1x final working concentration. The recommended loading phase for FLIPR Calcium 5 is 1hr, recording continued for 24hrs. Raw transient data for the windows indicated together with transient amplitude and beat rate over the first two hours of the experiment were extracted using the RTCA software and prepared for presentation using Microsoft Excel.

### Predictive structural modelling

The predicted models for the RGECO-TnT fusion were generated using RaptorX^44^and Swiss-PDB Viewer^45^ using the primary amino acid sequence provided in **Supplementary figure 12**.

## Author Contributions

PR and MJD conceptualized the study. PR, CR, MJD, AJS designed and interpreted the experiments. PR and AJS performed the chemical dye experiments. PR and AJS made the RGECO-troponin conjugate. AJS constructed and produced the RGECO-TnT adenovirus. PR, AJS, KS, MAG performed the characterisation of the GECO’s. CNB and AJS provided the RGECO-TnT predicted molecular model. CNB, Y-FC, FAB, XZ, YAA performed the iPS-CM experiments. PR and AJS performed the adult cardiomyocyte experiments. AJS provided analytic tools. AJS and PR performed all analysis. MJD, CR, PR, AJS wrote and edited the manuscript and all authors provided comments on this manuscript.

